# Hypoxia pathway proteins regulate the synthesis and release of epinephrine in the mouse adrenal gland

**DOI:** 10.1101/2020.10.15.340943

**Authors:** Deepika Watts, Nicole Bechmann, Ana Meneses, Ioanna K. Poutakidou, Denise Kaden, Catleen Conrad, Anja Krüger, Johanna Stein, Ali El-Armouche, Triantafyllos Chavakis, Graeme Eisenhofer, Mirko Peitzsch, Ben Wielockx

**Affiliations:** Institute of Clinical Chemistry and Laboratory Medicine, Technische Universität Dresden, 01307 Dresden, Germany; German Institute of Human Nutrition Potsdam-Rehbruecke, Department of Experimental Diabetology, 14558 Nuthetal, Germany; German Center for Diabetes Research (DZD), 85764 München-Neuherberg, Germany; Department of Pharmacology and Toxicology, Medical Faculty, Technische Universität Dresden, 01307 Dresden, Germany

**Author notes:** Correspondence to: Ben Wielockx, Institute of Clinical Chemistry and Laboratory Medicine, Technical University Dresden, Fetscherstrasse 74, 01307 Dresden, Germany.

## Abstract

The adrenal gland and its hormones regulate numerous fundamental biological processes; however, the impact of hypoxia signalling on its function remains scarcely understood. Here, we reveal that deficiency of HIF (Hypoxia Inducible Factors) prolyl hydroxylase domain protein-2 (PHD2) in the adrenal medulla of mice results in HIF2α-mediated reduction in phenylethanolamine N-methyltransferase (PNMT) expression, and consequent reduction in epinephrine synthesis. Concomitant loss of PHD2 in renal erythropoietin (EPO) producing cells stimulated HIF2α-driven EPO overproduction, excessive RBC formation (erythrocytosis) and systemic hypoglycaemia. Using mouse lines displaying only EPO-induced erythrocytosis or anaemia, we show that hypo- or hyperglycaemia is necessary and sufficient to respectively enhance or reduce exocytosis of epinephrine from the adrenal gland. Based on these results, we propose that the PHD2-HIF2α axis in the adrenal medulla and beyond regulates both synthesis and release of catecholamines, especially epinephrine. Our findings are also of great significance in view of the small molecule PHD inhibitors being tested in phase III global clinical development trials for use in renal anaemia patients.

## INTRODUCTION

The hypoxia signaling pathway regulates the expression of myriad genes involved in various biological processes in living animals, and the Hypoxia Inducible Factors (mainly HIF1 and 2) are the central transcription factors that regulate these processes. HIF expression, in turn, is under the direct control of a set of oxygen sensors known as the HIF-prolyl hydroxylase domain-containing proteins (PHD1-3). Under normoxic conditions, PHDs use oxygen as a co-factor to hydroxylate two prolyl residues in the HIFα subunits, thereby making HIFs accessible to the von Hippel-Lindau protein complex (pVHL) for subsequent ubiquitination and degradation [1]. Reduced cellular oxygen levels preclude such hydroxylation of HIFs by PHDs, resulting in stabilization of HIFα and direct transcriptional activation of more than 1000 genes. HIF transcriptional targets are primarily involved in a wide range of biological processes that serve to reverse the unfavourable hypoxic state, like erythropoiesis, blood pressure regulation and cell survival [2, 3]. Hypoxia is also a central feature of multiple pathologies, including local and systemic inflammation, and various stages of carcinogenesis, including pheochromocytomas, which are tumors originating from the adrenal chromaffin cells [4].

Adrenal chromaffin cells are part of the adrenal medulla and, as the source of catecholamines, are crucially involved in the fight-or-flight response, which requires epinephrine secretion [5]. Biosynthesis of epinephrine, an important catecholamine, is a complex multistep process in which the last step involves methylation of norepinephrine into epinephrine. This is catalysed by the enzyme phenylethanolamine N-methyltransferase (PNMT). Whereas all human chromaffin cells in the adrenal medulla produce epinephrine, in rodents, 20% of these cells don’t express PNMT and produce only norepinephrine [6]. Several *in vitro* studies have focused on HIF involvement in regulating the enzymatic activity required for catecholamine synthesis, and chromaffin cell-derived tumor cell lines have shown an essential role for HIF2α in catecholamine production; specifically, it regulated the expression of intermediate enzymes, namely, dopamine β-hydroxylase (DBH) and Dopa decarboxylase (DDC) [7]. Conversely, another *in vitro* study found no impact of HIF2α expression on tyrosine hydroxylase (TH) or Dbh [8], while downregulating PNMT expression [9]. The latter finding is in line with results from a number of other studies that have connected HIF2α activity in the adrenal medulla or in pheochomracytomas with reduced PNMT production [9-11]. Tumors with pVHL mutations also overexpress HIF2 target genes such as erythropoietin (EPO), a hormone central to red blood cell (RBC) formation [12, 13]. Despite these observations, studies focusing on how modulations of hypoxia pathway proteins affect adrenal function are rather sparse, and only recently has a transgenic mouse line harbouring a whole body HIF2α gain-of-function mutation been described, which showed reduced PNMT levels in the adrenal glands [14].

The concentration of epinephrine after stimulation of the sympathetic nervous system, including through hypoglycemia [5, 15] can increase up to 30-fold in circulation [16-18]. Even though hyperglycemia is associated with inhibition of insulin exocytosis from β-cells [19, 20], it is unknown if also influences epinephrine release from chromaffin cells. Interestingly, elevated levels of systemic EPO or treatment with erythropoiesis stimulating agents are directly linked to a reduction in blood glucose levels (BGL), possibly due to higher glucose consumption by greater numbers of RBCs [21-23]. As the production and release of epinephrine are critical, *in vivo* studies are essential to better understand the direct and/or indirect impact of alterations in hypoxia pathway proteins on epinephrine production and release from the adrenal gland.

Here, we performed an in-depth study of the synthesis of epinephrine and its release from the adrenal gland using several transgenic mice that exhibit functional changes in one or more hypoxia pathway proteins. Our results demonstrate that PHD2 deficiency in the adrenal medulla results in HIF2α-mediated reduction in PNMT and consequent inhibition of epinephrine synthesis in the adrenal gland. Moreover, we show that enhanced exocytosis of epinephrine into circulation is dependent on EPO-induced hypoglycemia.

## MATERIALS AND METHODS

### Mice

All mouse strains were maintained under specific pathogen-free conditions at the Experimental Centre of the Medical Theoretical Center (MTZ, Technical University of Dresden - University Hospital Carl-Gustav Carus) or at the animal facility of Max Planck Institute of Molecular Cell Biology and Genetics (MPI-CBG), Dresden. Experiments were performed with both male and female mice aged between 6-12 weeks. No significant differences between the genders were observed. CD68:cre-PHD2/HIF1^ff/ff^ (P2H1) or CD68:cre-PHD2^f/f^ (P2) lines were generated in-house and have been previously described by us [24]. CD68:cre-HIF1^f/f^ mice were generated for the current work using the HIF1^f/f^ line [25]. EPO Tg6 mice have been described earlier [26]. FOXD1:cre-HIF2α^f/f^ mice have been described elsewhere [27], but were generated in our laboratory using the FOXD1:cre line [28] (generous gift from Dr. Todorov, Dresden, Germany) in combination with the HIF2^f/f^ line [29]. All mice described in this report were born in normal Mendelian ratios. Mice were genotyped using primers described in supplementary Table 1. Peripheral blood was drawn from mice by retro-orbital sinus puncture using heparinized microhematocrit capillaries (VWR, Germany), and plasma separated and stored at −80°C until further analysis. Urine was immediately frozen on dry ice after collection and stored at −80°C until further analysis. Mice were sacrificed by cervical dislocation and adrenals were isolated, snap frozen in liquid nitrogen, and stored at −80°C for hormone analysis or gene expression analysis. All mice were bred and maintained in accordance with facility guidelines on animal welfare and with protocols approved by the Landesdirektion Sachsen, Germany.

### Laser microdissection

Adrenal glands were embedded in O.C.T Tissue-Tek (A. Hartenstein GmbH) and subsequently frozen on dry ice and stored at −80°C until further processing. Serial sections of 25-30 μm were obtained at −14°C using a cryotome. The glass slides used for collection of sections were heated at 200°C for a minimum of 3 hours and UV treated for 15 minutes. After collection of sections, slides were subjected to sequential dehydration in H_2_O, 75% EtOH, 95% EtOH and 100% EtOH for one minute each and then left to dry completely. Microdissection of the adrenal medulla was performed using a PALM MicroBeam LCM (ZEISS). Tubes containing the tissue were frozen to −20°C until further processing and genomic DNA was isolated from the dissected tissue using the alkaline lysis buffer followed by the neutralization buffer.

### Hormone measurement

Adrenal glands were incubated in disruption buffer (component of Invitrogen^TM^ Paris^TM^ Kit, AM 1921, ThermoFisher Scientific) for 15min at 4°C, homogenized in a tissue grinder, followed by incubation for 15 min on ice and further preparation. Adrenal *catecholamines*, norepinephrine, epinephrine, and dopamine were measured by high pressure liquid chromatography (HPLC) coupled with electrochemical detection after batch extraction with alumina, as previously described [30]. *Urinary catecholamines* were determined by liquid chromatography tandem mass spectrometry (LC-MS/MS) as described previously [31]. All urine samples were normalized for their volume by urinary creatinine measurement.

### PNMT enzyme activity

Phenylethanolamine N-methyltransferase (PNMT) enzyme activity was analyzed by LC-MS/MS as recently shown [32].

### RNA extraction and qPCRs

RNA from the adrenal glands was isolated using the RNA Easy Plus micro kit (Qiagen). cDNA synthesis was performed using the iScript cDNA Synthesis Kit (BIO-RAD). Gene expression levels were determined by quantitative real-time PCR using the ‘Ssofast Evagreen Supermix’ (BIO-RAD). Primer sequences used for qPCRs are included in supplementary Table 2. Expression levels of genes were determined using the Real-Time PCR Detection System-CFX384 (BIO-RAD). All mRNA expression levels were calculated relative to housekeeping genes, β2M or EF2, and were normalized using the ddCt method. Relative gene expression was calculated using the 2(-ddCt) method, where ddCT was calculated by subtracting the average WT dCT from dCT of individual samples.

### Cell culture

Mouse pheochromocytoma cells (MPC – 4/30/PRR) were obtained from Arthur Tischler (Department of Pathology and Laboratory Medicine, Tufts University School of Medicine, Boston, MA, USA; and Dr. Pacak, NIH, Bethesda, MD) [33]. The MPC cells were plated on collagen A-coated flasks (Biochrom AG, Berlin, Germany) and maintained in RPMI 1640 medium containing 5% foetal calf serum and 10% horse serum (all from Life Technologies, Darmstadt, Germany) at 37°C, 95% humidity, and 5% CO2.

### Intracellular calcium uptake

MPCs were cultured in 24-well plates in low serum medium – Opti-MEM (Thermo Fisher Scientific). Cells were plated onto collagen-coated plates in Opti-MEM for 3 hours and subjected to respective treatment conditions of either glucose (D-(+)-Glucose (Sigma Aldrich) or erythropoietin (Roche). Cells were harvested 24 hours later, washed, transferred to a 96-well plate and subjected to staining with 0.5 μmol Fluo-8-AM (Abcam) for 60 min at 37°C in assay bu er (HBSS with Pluronic F127 plus), followed by washing with HBSS and analysis on a BD FACS Canto.

### Statistical analyses

All data are presented as mean ± SEM. Data (WT control versus transgenic line) were analysed using the Mann–Whitney U-test, or the unpaired t-test with Welch’s correction, as appropriate (after testing for normality with the F test). All statistical analyses were performed on GraphPad Prism ver. 7.02 for Windows (GraphPad Software, La Jolla California USA, www.graphpad.com). Significance was set at p<0.05; ‘n’ in figure legends denotes individual samples.

## RESULTS

### Alterations in hypoxia pathway proteins reduce adrenal epinephrine

Previously, we have described a mouse line with conditional PHD2 and HIF1α inactivation in a variety of cells (CD68:cre-PHD2/HIF1α^ff/ff^ –henceforth designated P2H1), including in neurons and renal EPO producing cells (REPC), which results in local HIF2α stabilization that leads to excessive systemic EPO and erythrocytosis (Supplementary Figure 1A) [24]. As these cell lineages are thought to derive from neural crest cells [27, 34, 35], we looked at the effects of modulating hypoxia pathway proteins in another neural crest-derived cell type, namely, chromaffin cells, which are located in the medulla of the adrenal gland. First, genomic PCR on laser micro-dissected adrenal medullary tissue confirmed localized targeting of PHD2 and HIF1α (Supplementary Figure 1B), and qPCR analysis using mRNA from whole adrenal glands showed reduction of *Phd2* and *Hif1*α compared to the WT littermate controls, apart from an associated increase in *Hif2*α (Supplementary Figure 1C). Next, catecholamine levels in adrenal gland lysates showed dramatic decrease in only epinephrine in P2H1 mice compared to their WT littermates, but not of the upstream hormones, namely, dopamine and nor-epinephrine (Figure 1A). We also observed a corresponding marked decrease in PNMT mRNA, protein, and enzymatic activity (Figure 1B-D). Conversely, no differences in the expression of other catecholamine-associated enzymes, such as *Th* or *Dbh*, were detected (Supplementary Figure 1D). Taken together, these results show that inhibition of the oxygen sensor PHD2 and one of its downstream HIF targets in the adrenal gland leads to diminished epinephrine synthesis.

**Figure 1.**
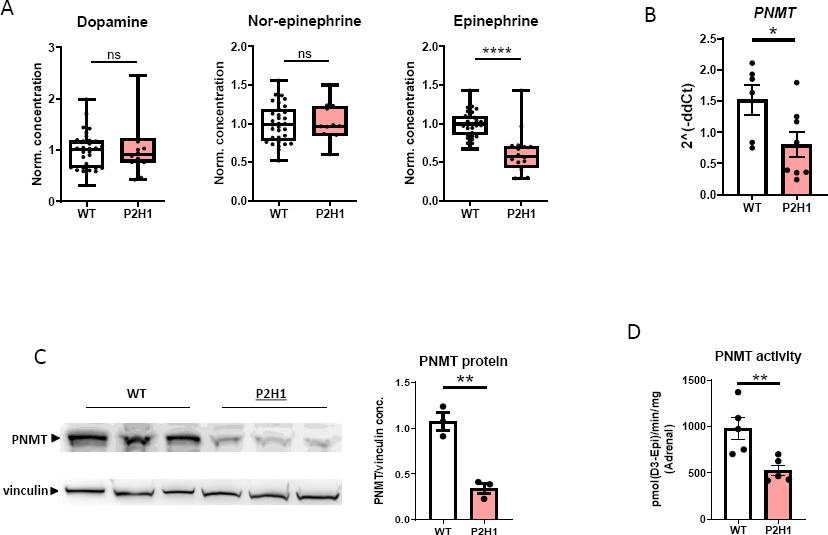
Loss of PHD2 and HIF1 results in decreased epinephrine synthesis and PNMT activity in the adrenal gland. A) Box and whisker plots showing catecholamine measurements in the adrenals from WT mice in comparison to littermate P2H1 mice (n=16-31 individual adrenal glands). All data is normalized to average measurements in WT mice. The graphs are a representative result of at least 3 independent experiments. B) qPCR-based mRNA expression analysis of PNMT in the adrenal gland from individual P2H1 mice and WT littermates (n=6-8 individual adrenals). C) Western blot analysis and comparison of PNMT protein from the adrenals of P2H1 mice and WT controls (n=6 vs 3 individual adrenals). D) PNMT enzyme activity in the adrenals of P2H1 mice compared to the WT littermate controls. Statistical significance was defined using the Mann-Whitney U test (*p<0.05; **p<0.005).

### Decreased adrenal epinephrine synthesis is unrelated to loss of HIF1a

To verify the impact of the individual hypoxia pathway proteins, we took advantage of the CD68:cre-PHD2^f/f^ mouse line (henceforth designated P2) [2, 24] and a newly created CD68:cre-HIF1α^f/f^ line (henceforth designated H1). Similar to our findings in P2H1 mice, P2 mice exhibited significantly lower levels of epinephrine in the adrenal glands compared to their WT littermates (Figure 2A), along with reduced *PNMT* mRNA and enzyme activity. (Figure 2B). *Th* and *Dbh* levels remained unaffected in P2 mice (Supplementary Figure 2). However, compared to WT mice, H1 mice showed no differences in any of hormones measured (Figure 2C), strongly suggesting that loss of HIF1α alone does not play a significant role in catecholamine production in the adrenal gland. Importantly, these results support our initial observation that PHD2 alters epinephrine synthesis, probably due to HIF2α stabilization.

**Figure 2.**
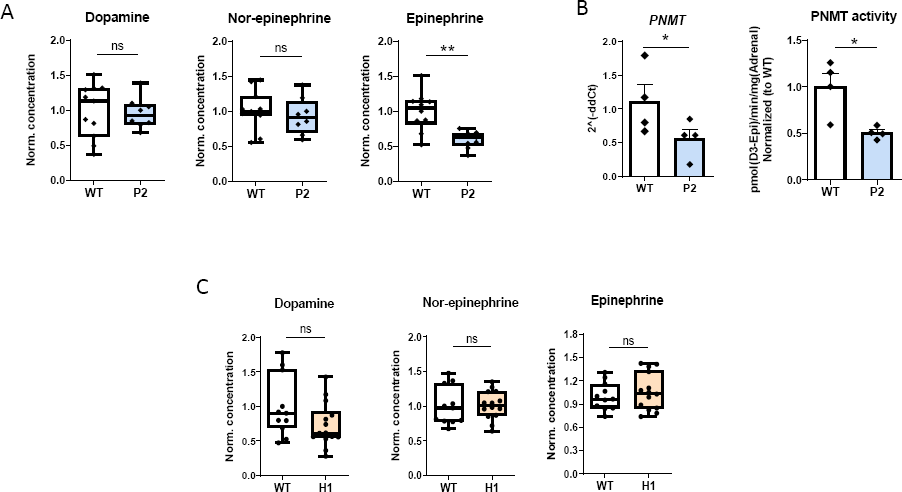
Decreased epinephrine and PNMT in the adrenal are related to the loss of PHD2 but unrelated to HIF1. A) Box and whisker plots comparing catecholamine measurements in the adrenals from P2 mice and WT littermates (n=8-11 individual adrenal glands). All data are normalized to average measurements in WT mice. The graphs are a representative result of at least 3 independent experiments. B) qPCR-based mRNA expression analysis of PNMT and PNMT activity in the adrenals of the individual mice (n= 4 vs 4). C) Catecholamine measurements in the adrenals of HIF1 mice compared to WT controls (n=11-14 individual adrenal glands). All data are normalized to the average of measurements in WT mice. Statistical significance was defined using the Mann-Whitney U test (*p<0.05; **p<0.005).

### Increased EPO is associated with PNMT-independent reduction in adrenal epinephrine synthesis

We have previously shown that both P2 and P2H1 mice, but not H1 mice, exhibit EPO-induced erythrocytosis [24] (Supplementary Figure 3A), and we wanted to investigate any potential relationship between reduced epinephrine levels in the adrenal glands of these mice and EPO-associated changes. Therefore, we tested the EPO transgenic mouse line (EPO Tg6) [36], which are known to constitutively produce high levels of RBCs without influencing alterations in adrenal hypoxia pathway proteins (Supplementary Figure 3A-B). Interestingly, similar to P2 and P2H1 mice, epinephrine levels were significantly lower in the adrenal glands of EPO Tg6 mice, along with dopamine levels (Figure 3A). In contrast to P2H1 and P2 mice, however, no differences in PNMT activity were observed, despite increased *PNMT* mRNA levels (Figure 3B). Congruently, mRNA analysis on whole adrenals revealed no changes in *Th* or *Dbh* (Supplementary Figure 3B). Taken together, these observations suggest that PNMT-independent reduction in adrenal epinephrine levels correlated to high systemic EPO.

**Figure 3.**
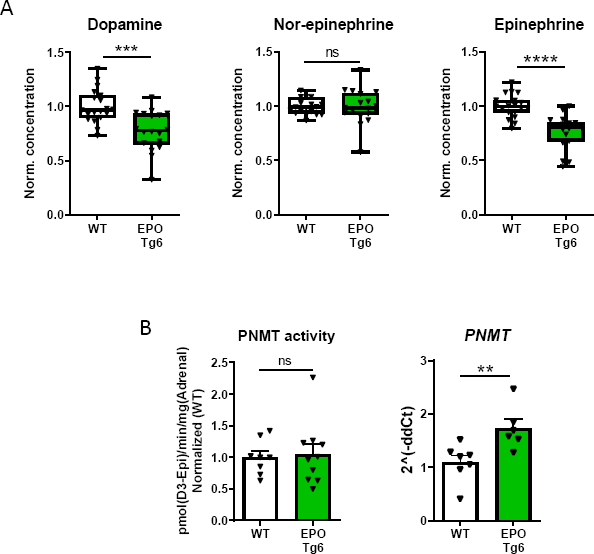
Lower epinephrine in the adrenals of EPO Tg6 mice. A) Box and whisker plots comparing catecholamine measurements in the adrenals from EPO Tg6 mice and WT littermates (n=19 vs 18 individual adrenal glands). All data are normalized to average measurements in WT mice. The graphs are a representative result of at least 3 independent experiments. B) qPCR-based mRNA expression analysis of PNMT and PNMT enzyme activity assay in the adrenals of individual mice (n= 8 vs 10). All data are normalized to the average of measurements in WT mice. Statistical significance was defined using the Mann-Whitney U test (*p<0.05; **p<0.005).

### EPO/erythropoiesis regulates epinephrine release from the adrenal gland

To characterize the impact of systemic EPO/erythropoiesis on epinephrine in adrenal glands, we looked at potential changes in secretion by measuring its levels in urine and also by calculating the ratio of epinephrine and its precursor, nor-epinephrine, i.e., EPI/NEPI. Interestingly, we detected a marked increase in both urinary epinephrine levels and in EPI/NEPI ratio in EPO Tg6 mice (Figure 4A), suggesting either direct or indirect effect of EPO on epinephrine release from the adrenal gland. Similarly, and despite PNMT-dependent inhibition of epinephrine in the adrenal gland (Figure 1A-B), P2H1 mice showed an almost significant increase in urinal epinephrine compared to WT littermates (Figure 4B).

**Figure 4.**
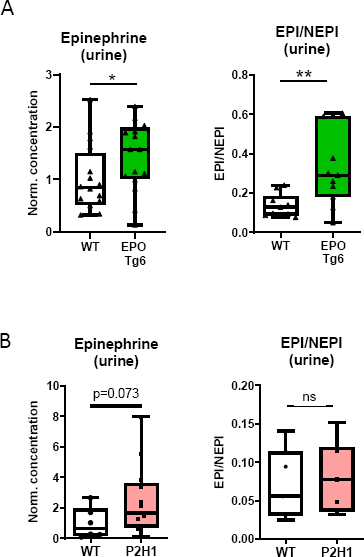
Enhanced urinary epinephrine and EPI/NEPI ratios in erythrocytotic Tg6 mice. A) Box and whisker plots comparing urinary Epinephrine (left) and EPI/NEPI ratios (right) in EPO Tg6 mice and WT littermates (n = 16 vs 16 mice). The graphs are a representative result of at least 3 independent experiments. B) Box and whisker plots comparing urinary epinephrine (left) and EPI/NEPI ratios (right) in P2H1 mice and WT littermates (n = 6 vs 13 mice). All urine measurements were normalized to urinary creatinine and data was further normalized to average measurements in WT mice. Statistical significance was defined using the one-tailed or two-tailed Mann-Whitney U test (*p<0.05; **p<0.005).

To further investigate these findings, we used a previously described anaemic mouse strain, namely, FOXD1:cre-HIF2α^f/f^ [27]. These mice show low systemic EPO levels and a dramatic reduction in circulatory RBCs (Supplementary Figure 4A) [27]. In direct contradiction to urinary data of the erythrocytotic EPO Tg6 mice, FOXD1:cre-HIF2α^f/f^ mice showed a >8-fold reduction in urinary epinephrine compared to WT littermates (Figure 5A). Interestingly, the adrenal glands of these anaemic mice showed a significant increase in dopamine, while levels of all other catecholamines remained unchanged (Figure 5B). Levels of neither PNMT (Figure 5C), nor of any of the other corresponding enzymes were altered (Supplementary Figure 4B). Taken together, at this point, our data strongly point towards EPO-dependent exocytosis of epinephrine that is independent of its synthesis in the adrenal.

**Figure 5.**
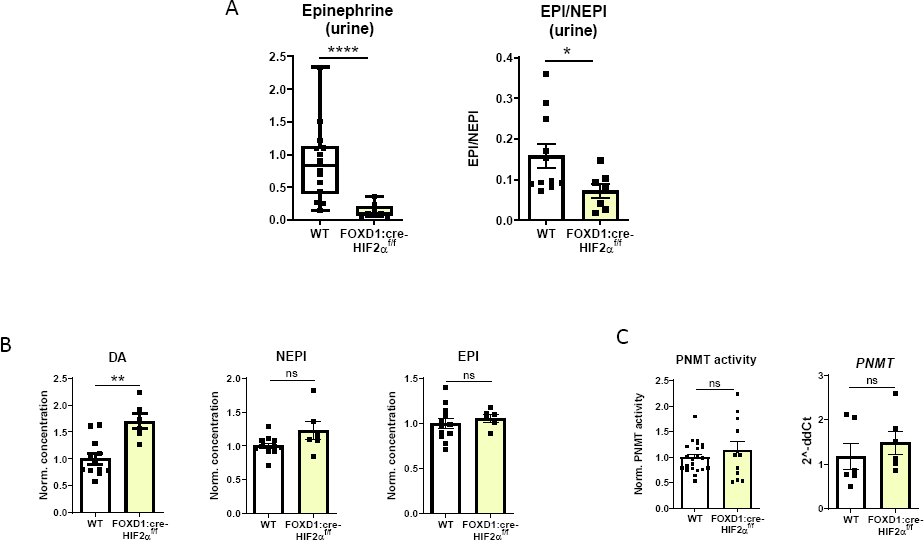
Decreased epinephrine and EPI/NEPI ratios in the urine of FOXD1:cre-HIF2α^f/f^ mice. A) Box and whisker plots comparing urinary epinephrine (left) and EPI/NEPI ratios (right) in FOXD1:cre-HIF2α^f/f^ mice and WT littermates (n = 14 vs 9 mice). The graphs are a representative result of at least 3 independent experiments. All urine measurements were normalized to urinary creatinine or urea levels and data was further normalized to average measurements in WT mice. B) Catecholamine measurements in the adrenals from FOXD1:cre-HIF2α^f/f^ mice compared to WT littermates (n=12 vs 6 individual adrenal glands). All data are normalized to the average measurements in WT mice. All urine measurements were corrected for differences in urine volumes by creatinine or urea concentrations in the urine. Data were further normalized to the average measurements in WT mice. The graphs are a representative result of at least 2 independent experiments. C) qPCR-based mRNA expression analysis of PNMT and PNMT enzymatic activity measurements in the adrenal of individual mice (n= 8 vs 10). All data are normalized to average measurements in WT mice. Statistical significance was defined using the Mann-Whitney U test (*p<0.05; **p<0.005).

### EPO-induced hypoglycaemia activates epinephrine release

Catecholamine release from chromaffin cells requires their exocytosis and this process is selectively stimulated by hypoglycaemia [18]. Recently, it has been shown that hyperactive erythropoiesis, as seen in EPO Tg6 mice, increases systemic glucose consumption and consequent hypoglycaemia, which is most probably attributable to a greater number of circulating RBCs [22]. Therefore, we sought to understand if and how EPO-associated hypoglycaemia can affect adrenal epinephrine release. We first measured blood glucose levels (BGL) in all mouse lines and found that both erythrocytotic P2H1 and EPO Tg6 mice display significant hypoglycaemia (Figure 6A), while conversely, the anaemic FOXD1:cre-HIF2α^f/f^ mice were dramatically hyperglycaemic (Figure 6B). Next, we used a mouse pheochromocytoma cell line (MPC) to study how changes in blood glucose levels could affect cellular exocytosis. That exocytosis is a calcium-dependent process is well-established, and increased calcium uptake by a cell is directly correlated with greater exocytosis [37, 38]; thus, calcium uptake is an appropriate readout of exocytosis. Next, we exposed MPCs to increasing concentrations of glucose and observed significant inhibition of calcium uptake, indicating diminished exocytosis (Figure 6C). In contrast, direct exposure of MPCs to high EPO did not lead to changes in intracellular calcium (Supplementary Figure 5A and B), clearly suggesting that EPO would not directly influence epinephrine exocytosis; rather it is systemic hypoglycaemia secondary to EPO/erythrocytosis in mice that enhances adrenal epinephrine release.

**Figure 6.**
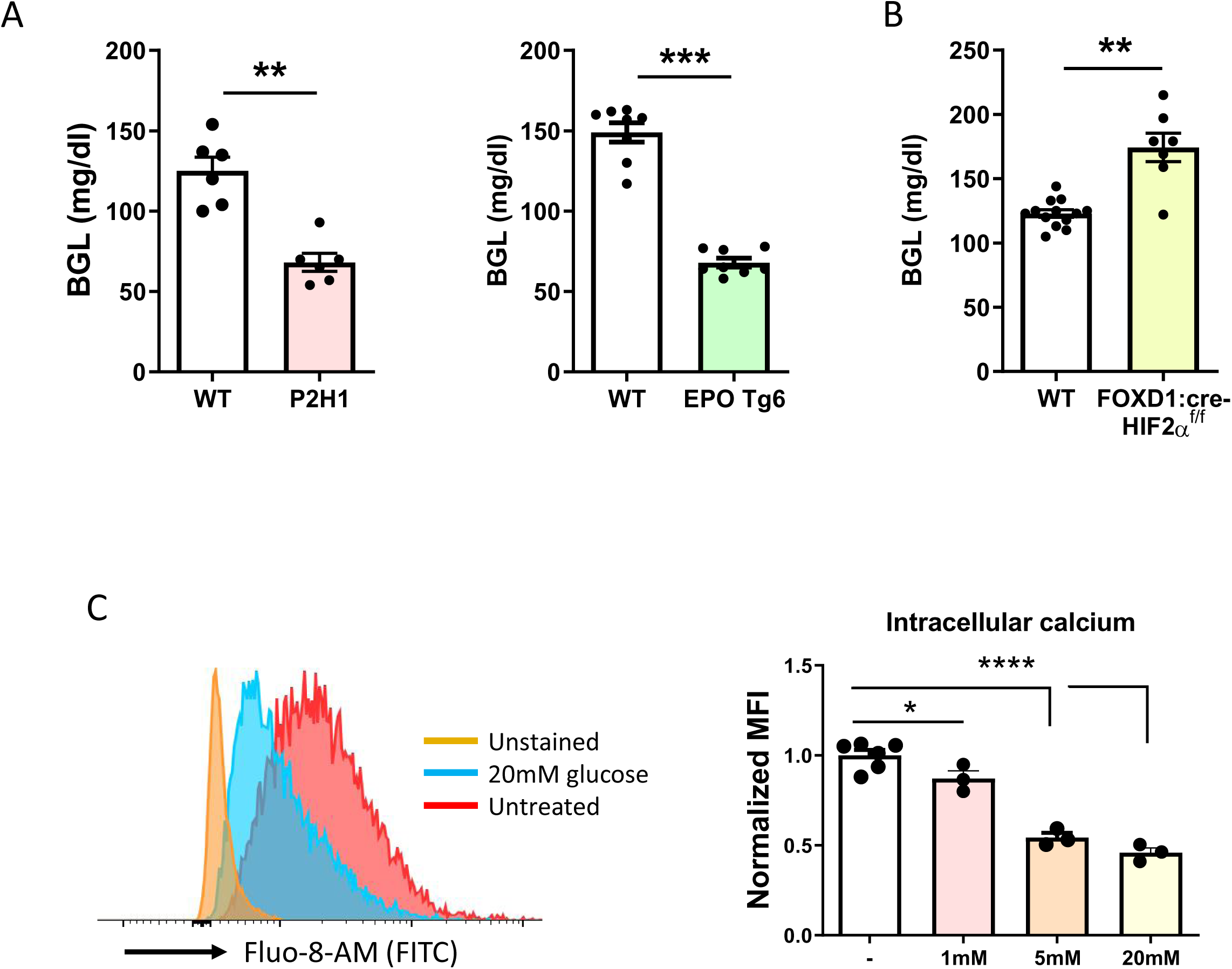
EPO-mediated regulation of blood glucose levels in erythrocytotic and anaemic mice. A) Blood glucose level (BGL) measurements in P2H1 and EPO Tg6 mice compared to WT littermates (n = 6-8 mice). B) BGL in FOXD1:cre-HIF2α^f/f^ mice compared to WT littermates (n = 6-13 mice). C) Representative histograms showing fluorescence of the Fluo-8-Am dye for intracellular calcium measurement in untreated MPC cells compared to stimulation with glucose. Normalized MFI depicting intracellular calcium measurement in MPC cells that were treated with glucose in a dose dependent manner. Each dot represents an individual well. The data is representative of 2 individual experiments. Statistical significance was defined using the Mann-Whitney U test or unpaired t-test (*p<0.05; **p<0.005).

## Discussion

Here, we show that decreased PNMT expression in the PHD2 deficient adrenal medulla reduces epinephrine synthesis, but systemic effects in these PHD2 deficient mice, such as enhanced EPO-production, consequent RBC excess, and hypoglycaemia in REPCs, lead to greater exocytosis, i.e., enhanced release of this hormone from the adrenal gland (Figure 7). These results indicate uncoupling between synthesis and excretion of epinephrine upon PHD2 loss. Mechanistically, we show that excessive epinephrine secretion is related to EPO-induced systemic hypoglycaemia, rather than a direct cell-level effect of EPO per se.

**Figure 7.**
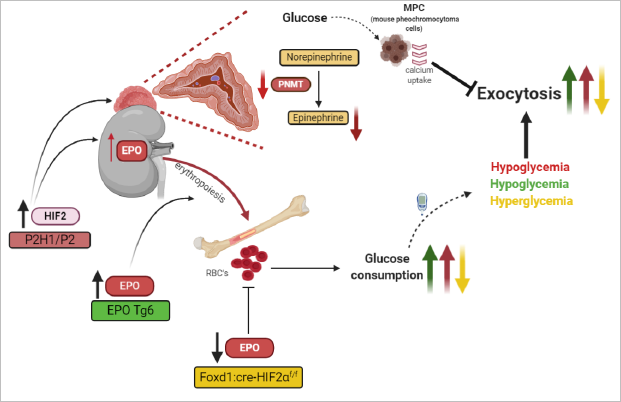
Graphical overview of the PHD2-HIF2α axis involved in synthesis and secretion of epinephrine from the mouse adrenal gland

Multiple hypoxia pathway components have been suggested to define the functionality of the adrenal medulla. In the physiological sympathoadrenal setting, pVHL has been shown to be essential during development and it is required by the peripheral oxygen-sensing system to ensure survival under hypoxic conditions [39]. Loss-of-function mutations in the *VHL* gene or gain-of-function mutations in *HIF2*α have been associated with the development of pheochromocytoma and paraganglioma [40]. However, this HIF-subunit is not essential for the development and functionality of the adrenal medulla, as mice deficient for HIF2α in TH-positive medullary cells display no functional abnormalities [41]. In contrast, a novel mouse line containing a HIF2α gain-of-function mutation has been reported to display reduced adrenal PNMT and metanephrine/normethanephrine ratios, although no data on potential modulation of catecholamines in the adrenal gland were provided [14]. Our data on HIF2α-induced reduction in PNMT activity concur with these observations. Likewise, several *in vitro* studies have described the effects HIF2α on the various enzymes involved in the process of catecholamine production in chromaffin-related tumour cell lines [7-9, 42, 43]. In contrast, the role of HIF1α in the adrenal medulla remains largely unexplored, although a recent study has linked the loss of HIF1α in the progenitors of the second heart field and the pharyngeal mesoderm to lower numbers of TH^+^ chromaffin cells in the adrenal gland [44]. Therefore, further insight into how PHDs/HIFs regulate *in vivo* epinephrine production and release from the adrenal medulla is essential, as there is a promising future for HIF stabilizers (HIF-PHD inhibitors) in the treatment of anaemia. However, it must be noted here that concerns have been recently raised regarding the chronic use of such HIF stabilizers [3, 45-48].

The advantage of our approach lies in the use of multiple transgenic mice lines to define the individual roles of PHD2 and HIF1α in epinephrine physiology, which was complimented by studies in mice that are conditionally deficient for both PHD2 and HIF1α in cells of neural origin i.e., REPCs [24, 49]. Observations from all these mouse lines strongly suggest targeting of chromaffin cells in the adrenal medulla. We have previously reported HIF2α stabilization in P2H1 mice and demonstrated that the excessive erythrocytosis phenotype seen in these P2H1 mice is attributable to HIF2α stabilization in the absence of the protective effects of HIF1α [24]. Similarly, we show that loss of PHD2 alone, but not of HIF1 alone, resembled the P2H1 phenotype with respect to changes in epinephrine synthesis and secretion, indicating that this phenotype too is enhanced through stabilization of HIF2α and that HIF1α might not have a significant role. Further, although we found a comparable decrease in epinephrine levels in P2H1 and P2 adrenal glands and a corresponding reduction in adrenal PNMT activity, urinary excretion of epinephrine was higher in P2H1 mice, suggesting that epinephrine was preferentially released from the adrenal glands. As P2H1 mice also display excessive erythrocytosis that is associated with very high levels of EPO, we hypothesized that this excessive EPO was responsible for the observed preferential release of epi. Therefore, we used erythrocytotic EPO Tg6 mice to understand the effects of only the EPO/polycythaemia phenotype on epinephrine physiology without interference from the HIF2-PNMT pathway [36], and show that excessive EPO indeed increases urinary epinephrine excretion and EPI/NEPI levels, thereby supporting our hypothesis.

Under basal conditions, exocytosis from the chromaffin cells is a strictly controlled process that largely depends on calcium uptake [50-53]. However, it can be further enhanced by additional stimuli, including hypoglycaemia, hypotension, hypoxemia and emotional distress [4, 17, 53, 54]. Interestingly, it has been recently reported that EPO Tg6 mice are strongly hypoglycaemic and that low BGL correlates with the degree of erythropoiesis [22, 55]. Moreover, it was postulated almost a century ago that hypoglycaemia can *per se* stimulate epinephrine release from the adrenal medulla [56]. The results of our in vitro experiments in MPCs lend support to this hypothesis as we show for the first time that calcium uptake reduces as glucose concentration increases, and that direct EPO exposure does not produce such an effect. Thus, based on our observations, we suggest, for the first time, that EPO-induced RBC excess, and consequent hypoglycaemia, lead to enhanced exocytosis of epinephrine that is mediated by enhanced Ca^2+^uptake in chromaffin cells. Further, higher PNMT mRNA and reduced dopamine levels in EPO Tg6 adrenal glands are most likely the consequence of such enhanced exocytosis. These conclusions are borne out by the contrasting observations in the anaemic FOXD1:cre-HIF2α^f/f^ mice [27] which show dramatically low levels of epinephrine in urine that were linked to hyperglycaemia, even though adrenal epinephrine and PNMT levels remained unchanged. We postulate that higher dopamine levels in these mice serve to avoid NEPI and EPI overproduction in the adrenal gland.

In summary, we show that PHD2-mediated HIF2α stabilization in the adrenal gland has divergent local and systemic effects *in vivo*, i.e., while epinephrine synthesis is diminished, its urinary excretion is enhanced. Importantly, in these mice, adrenal synthesis of epinephrine and its urinary secretion appear to be clearly uncoupled as excessive urinary epinephrine secretion appears to be due to systemic EPO-induced effects via the PHD2-HIF2-EPO axis, which include excessive erythrocytosis and consequent increase in glucose consumption by these RBCs. Mechanistically, we propose that the EPO-induced RBC excess and consequent hypoglycaemia lead to enhanced exocytosis of epinephrine by increasing Ca^2+^uptake in chromaffin cells, irrespective of the primary reduction in adrenal epinephrine synthesis. These findings have potential clinical significance in view of broad spectrum HIF-PHD inhibitors currently under Phase III trials.

## Acknowledgments

This work was supported by grants from the DFG (German Research Foundation) within the CRC/Transregio 205/1, Project No. 314061271 - TRR205, “The Adrenal: Central Relay in Health and Disease” (A02) to B.W., T.C, A-E-A.; (B12) to N.B., G.E; (S01) to D.K., M.P.; B.W. was supported by the Heisenberg program, DFG, Germany; WI3291/5-1 and 12-1). We would like to thank Dr. Vasuprada Iyengar for English Language and content editing.

## Author contributions

D.W. designed and performed a majority of the experiments, analysed data, and wrote the manuscript. N.B. performed experiments, analysed data, and contributed to the discussion. A. Me. provided essential tools, performed experiments and analysed data. D.K., A.K., J.S. performed experiments and analysed data. A.E.A. and T.C. provided tools and contributed to the data discussion. G.E. and M.P. provided tools, analysed data and contributed to the data discussion. B.W. designed and supervised the overall study, analysed data, and wrote the manuscript.

## Supplementary Figures

**Supplementary figure 1. Blood and adrenal analysis of P2H1 mice for HCT/HGB data and gene expression, respectively.**

A) Bar graphs showing increased RBC, HCT and HGB in the blood of P2H1 mice. B) Genomic PCR analysis of laser microdissected adrenal medulla samples from WT and P2H1 mice. C) Bar graphs showing the qPCR-based mRNA expression analysis of PHD2, HIF1 and HIF2 from the adrenals (n=6-7) of P2H1 mice and littermate WT controls. D) Bar graphs showing the qPCR-based mRNA expression analysis of Th and Dbh from the adrenals (n=6-7) of P2H1 mice and the littermate WT controls. All the data are normalized to average measurements in WT mice. Statistical significance was defined using one-tailed or two-tailed Mann-Whitney U test, as required (*p<0.05; **p<0.005).

**Supplementary figure 2. Gene expression analysis in the adrenals of P2 mice.**

Bar graphs showing the qPCR-based mRNA expression analysis of Th and Dbh from the adrenals (n=4 vs 4) of the P2 mice and littermate WT controls. All the data are normalized to the average measurements in WT mice.

**Supplementary figure 3. Blood analysis for HCT/HGB data and gene expression analysis in the adrenals of EPO Tg6 mice.**

A) Bar graphs showing increased RBC, HCT and HGB in the blood of EPO Tg6 mice. B) Genomic PCR analysis on laser microdissected adrenal medulla samples from WT and P2H1 mice. C) Bar graphs showing the qPCR-based mRNA expression analysis of PHD2, HIF1 and HIF2 from the adrenals (n=6-7) of EPO Tg6 mice and littermate WT controls. D) Bar graphs showing qPCR-based mRNA expression analysis of *Th* and *Dbh* from the adrenals (n=6-7) of EPO Tg6 mice and littermate WT controls. All the data are normalized to average measurements in WT mice. Statistical significance was defined using the Mann-Whitney U test (*p<0.05; **p<0.005).

**Supplementary figure 4. Blood analysis for HCT/HGB and gene expression analysis in the adrenals of FOXD1:cre-HIF2α^f/f^ mice.**

A) Bar graphs showing increased RBC, HCT and HGB in the blood of FOXD1:cre-HIF2α^f/f^ mice. B) Bar graphs showing the qPCR-based mRNA expression analysis of *Th* and *Dbh* from the adrenals (n=6 vs 6) of FOXD1:cre-HIF2α^f/f^ mice and littermate WT controls. All the data are normalized to the average of measurements in WT mice. Statistical significance was defined using the Mann-Whitney U test (*p<0.05; **p<0.005).

**Supplementary figure 5: Intracellular calcium measurement in the EPO-treated MPC cells.**

A) Representative histograms showing fluorescence of Fluo-8-Am dye, depicting intracellular calcium in the untreated MPC cells compared to treatment with EPO. B) Normalized MFI depicting intracellular calcium measurement in EPO-treated MPC cells. Each dot represents an individual well. The data is representative of 2 individual experiments. Statistical significance was defined using the Mann-Whitney U test (*p<0.05; **p<0.005).

