## Supplemental Figures for "Hypoxia pathway proteins regulate the synthesis and release of epinephrine in the mouse adrenal gland"

Supplementary Figure 1

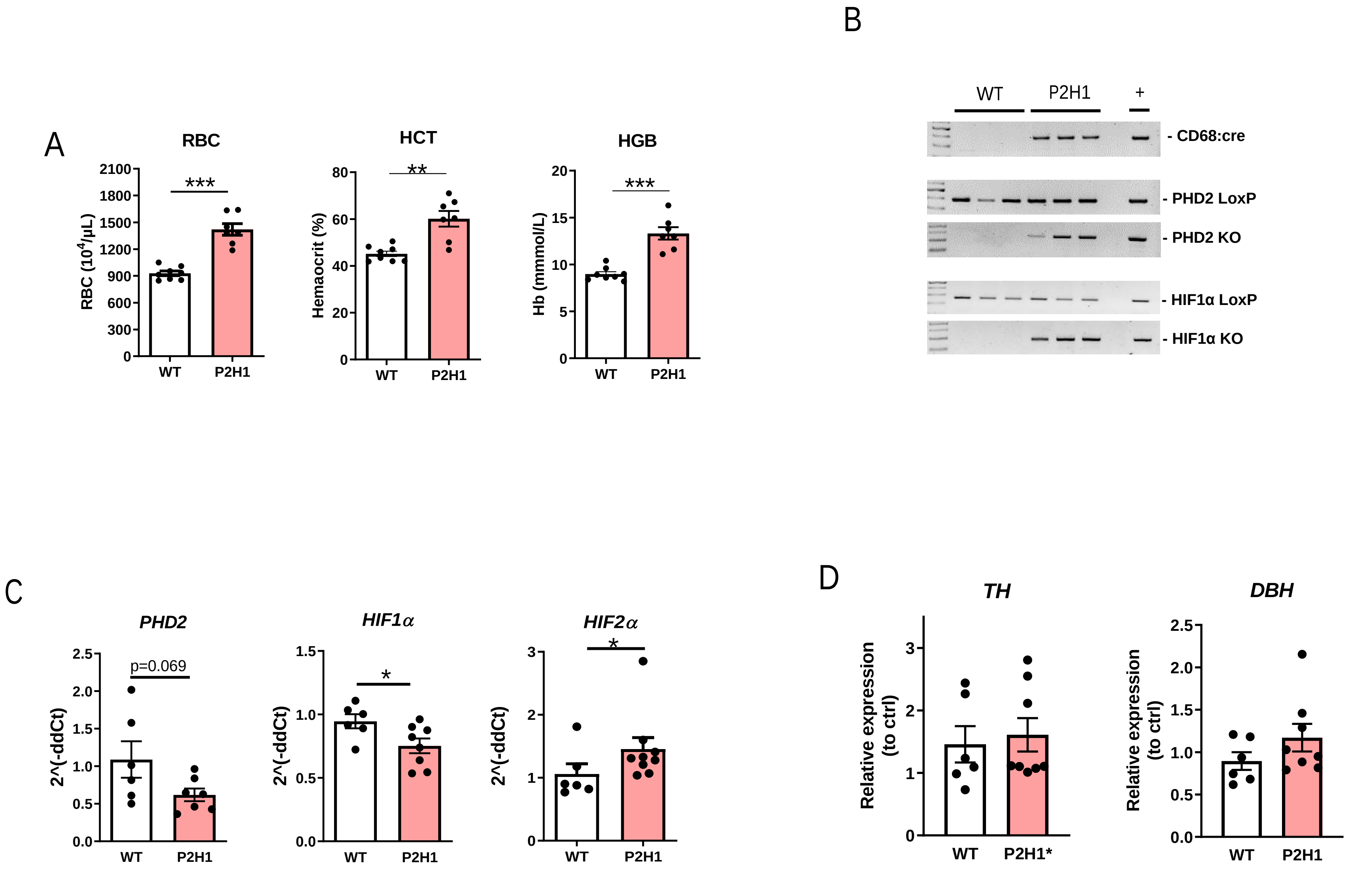

Supplementary Figure 2

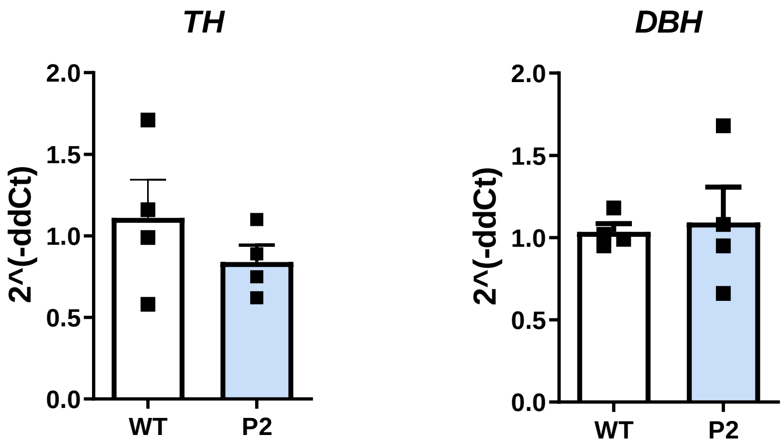

Supplementary Figure 3

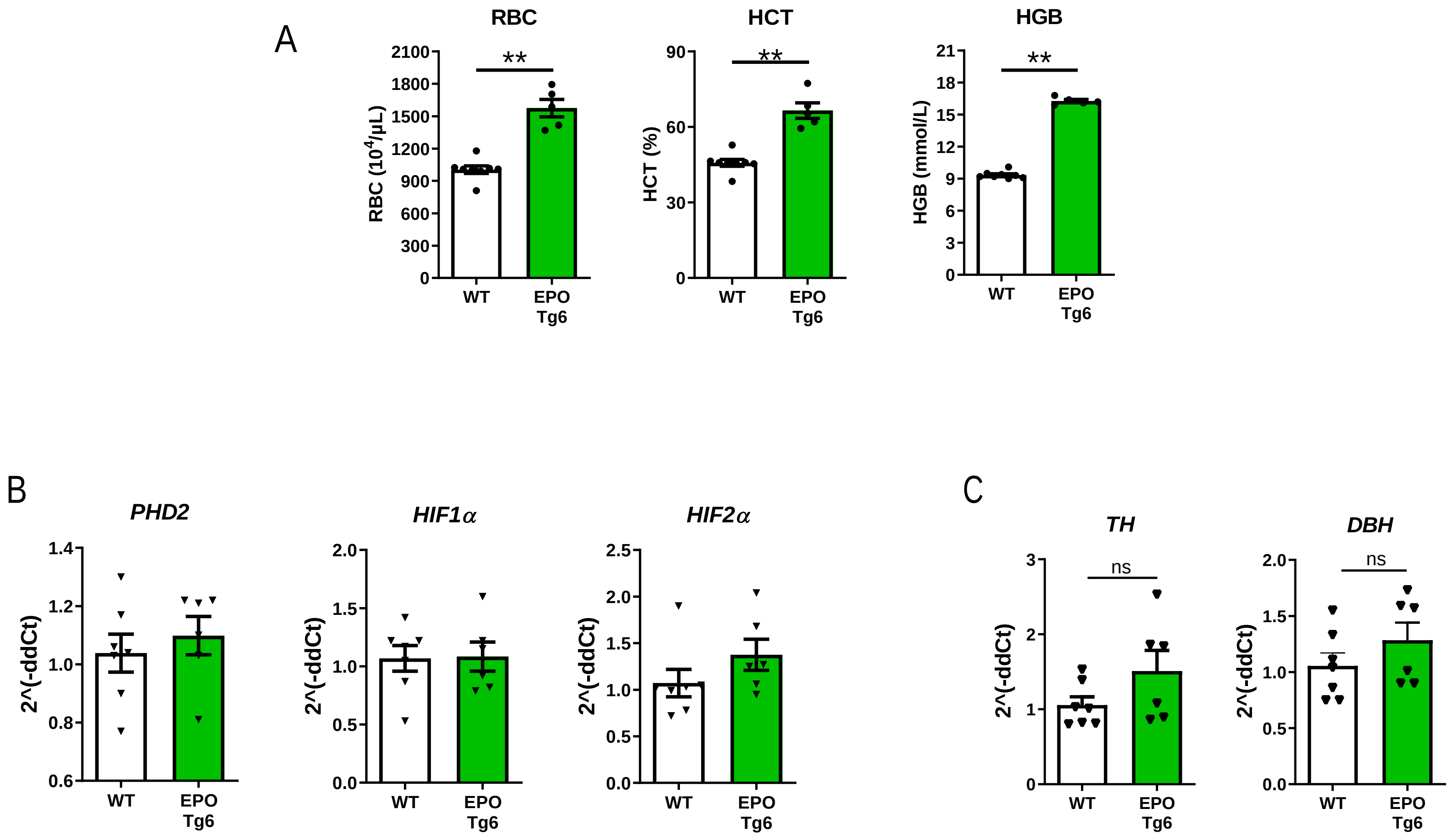

Supplementary Figure 4

A

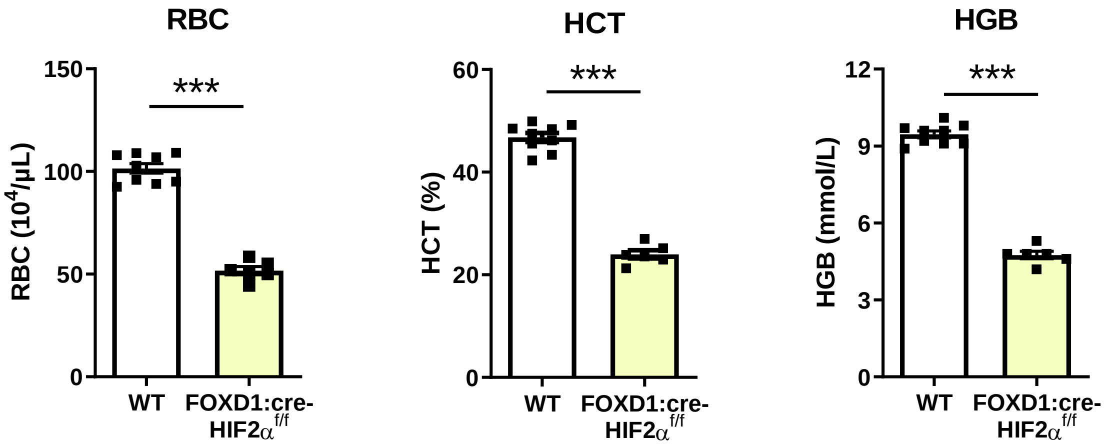


B

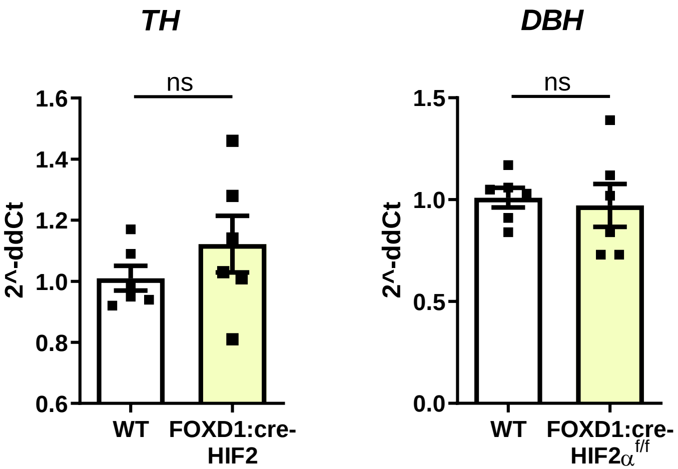

Supplementary Figure 5

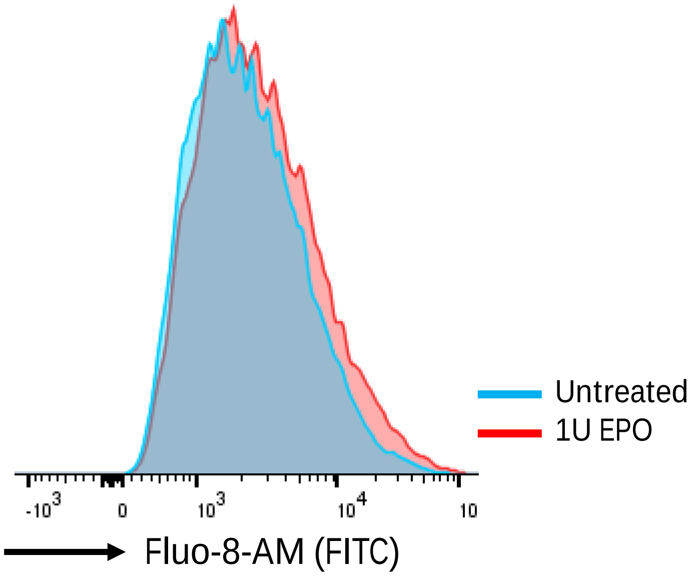


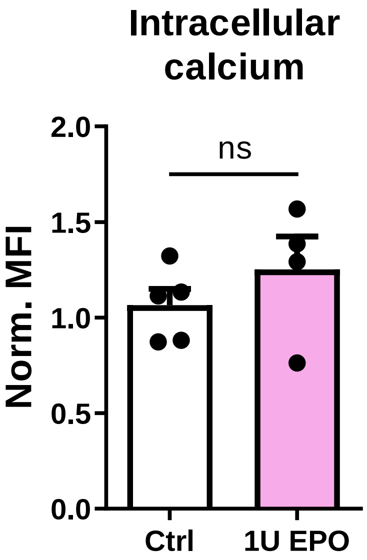
