## Supplemental Data for "Hypoxia pathway proteins regulate the synthesis and release of epinephrine in the mouse adrenal gland"

Table I : Primers for genotyping of mouse strains

| **Primer name** | **Primer sequence (5’ – 3’)** |
| --- | --- |
| Cre_Fwd  Cre_Rev | GCCTGCATTACCGGTCGATGCAACGA  GTGGCAGATCGCGCGGCAACACCATT |
| HIF1α_flox_Fwd  HIF1α_flox_Rev | GCAGTTAAGAGCACTAGTTG  GGAGCTATCTCTCTAGACC |
| HIF-1α_AR1115 | GCTACTGTAAATTTGGGGATGAAAACATCTGCT |
| HIF-1α_AR1116 | GCAGTTAAGAGCACTAGTTGATCTTTCCGAGG |
| mPHD2_exo2  mPHD2_Intron1 | CGCATCTTCCATCTCCATTT  CTCACTGACCTACGCCGTGT |
| mPHD2_Intron1  mPHD2_Intron3.3 | CTCACTGACCTACGCCGTGT  GGCAGTGATAACAGGTGCAA |

Table II: Primers for qPCR analysis

| **Primer name** | **Primer sequence (5’ – 3’)** |
| --- | --- |
| HIF1α_Fwd  HIF1α_Rev | GGCGAGAACGAGAAGAAAAA  AAGTGGCAACTGATGAGCAA |
| mPHD2_Fwd  mPHD2_Rev | AAGCCCAGTTTGCTGACATT  CTCGCTCATCTGCATCAAAA |
| HIF2α_Fwd  HIF2α_Rev | CTGAGGAAGGAGAAATCCCGT  TGTGTCCGAAGGAAGCTGATG |
| TH_Fwd  TH_Rev | TACAAGCAGGGTGAGCCAAT  TGGGTAGCATAGAGGCCCTT |
| DBH_Fwd  DBH_Rev | TGGGTGCCAAGGCATTTTAC  TGTGTAGTGTAGGCGGATGC |
| PNMT_Fwd  PNMT_Rev | CGCCTATCTCCGCAACAACT  GACACCTCACCGGTAGCAAAG |
